# Organisation of the transcriptional regulation of genes involved in protein transactions in yeast

**DOI:** 10.1101/229039

**Authors:** Duygu Dikicioglu, Daniel J H Nightingale, Valerie Wood, Kathryn S Lilley, Stephen G Oliver

## Abstract

The topological analyses of many large-scale molecular interaction networks often provide only limited insights into network function or evolution. In this paper, we argue that the functional heterogeneity of network components, rather than network size, is the main factor limiting the utility of topological analysis of large cellular networks. We have analysed large epistatic, functional, and transcriptional regulatory networks of genes that were attributed to the following biological process groupings: protein transactions, gene expression, cell cycle, and small molecule metabolism. Control analyses were performed on networks of randomly selected genes. We identified novel biological features emerging from the analysis of functionally homogenous biological networks irrespective of their size. In particular, direct regulation by transcription as an underrepresented feature of protein transactions. The analysis also demonstrated that the regulation of the genes involved in protein transactions at the transcriptional level was orchestrated by only a small number of regulators. Quantitative proteomic analysis of nuclear- and chromatin-enriched sub-cellular fractions of yeast provided supportive evidence for the conclusions generated by network analyses.

## Background

Biological activities at the cellular level are not isolated but are governed by a set of concerted interactions between the functional molecules responsible for performing a given task. For this reason, network analysis is often employed to elucidate how these coordinated interactions facilitate the functioning of an entire biological system (1). The connectivity graph of a cellular network, in which biological entities (such as genes, proteins, or metabolites) are connected through edges that represent interactions (such as reactions or structural associations) is often employed to identify the network’s topological features (2,3). Examples of the theoretical analysis of such graphs were reported by Erdös and Rényi as early as the 1960s, in a study of large random networks (4).

In early studies of biological networks, the topological structures were observed to depart from those of random networks. Biological networks were found to exhibit the following characteristics: a power law degree distribution, scale-freeness, ‘small-world’ properties, high tolerance to random attacks and low tolerance to targeted attacks, growth by preferential attachment, and the exhibition of distinct motif signatures and dense modules (5-8). Deviations from these characteristics (7) were usually attributed to incomplete or poor-quality data (6,9,10). However, such “exceptions” persisted, even after high-quality interaction data became available and advanced network analysis procedures came into play. This required the modification and adaptation of existing models of biological network topology, with many of their classical features being challenged as mere artefacts inherent in the representation of complex molecular interactions as two-dimensional network graphs (5). This has resulted in biological network analysis becoming increasingly focused, not on the global network, but on the identification of functionally relevant sub-structures within the network (5,9,11). The identification of tightly interconnected substructures (or modules) has proved to be a valuable tool with which to explore the workings of living cells in both control/healthy or modified/diseased states (12-14). Moreover, the study of even smaller local substructures of only a few nodes has allowed the identification of network motifs that have predictive value for interactions of similar nodes in different organisms or systems (15).

In this paper, we explore the importance of functional homogeneity in gaining biological insights from the topological analysis of cellular networks. Using the yeast *Saccharomyces cerevisiae* as our model, we have compiled sets of genes, each of which could be attributed to a broad biological process: e.g. the biosynthesis, modification, localization, and degradation of proteins (which we define here as protein transactions); gene expression (including both transcription and translation); cell cycle; or small molecule metabolism. We then constructed functional, regulatory, or epistatic interaction networks for these gene sets, and investigated their topological properties. We observed that the large size of the networks under investigation was not the major factor limiting the power of these analyses, and that it was possible to gain biological insights from the global topological analysis of these functionally homogenous networks. We then focused on the distinctive transcriptional regulatory patterns governing protein transactions in yeast, and carried out an indepth investigation of the nuclear proteome to confirm the inferences from our global topological analysis.

## Methods

### The gene and annotation datasets

Seven different gene product sets were investigated in the study. Gene Ontology (GO) Biological Process terms were employed to compile the function-specific sets (16) of *Saccharomyces cerevisiae* gene products associated with the following GO IDs: (i) protein transactions (PT) (G0:0006457, G0:0008104, G0:0019538, G0:0006986), (ii) gene expression (GE) (G0:0010467), (iii) cell cycle (CC) (G0:0007049), and (iv) small molecule metabolism (SMM) (G0:0044281). The GO term association data for these four sets and for the global set (Biological Process root node GO: 0008150) (6447 genes) were accessed from http://www.geneontology.org/ (access date 03/03/2017) (17). A pool of 1000 random sets, with equal size to that of PT (2533 gene products), was constructed by randomly selecting entries from the global set. One of these random sets was, itself, randomly selected and used for further analyses. The PT-C (complement of PT) set was constructed from all the non-PT gene products of the global set (3914). Signalling-associated gene products were identified using the GO signalling term (G0:0023046). Haplosufficiency, essentiality, Gene Ontology associations, and human protein functional homology information was retrieved from the Saccharomyces Genome Database accessed on 22/03/2017 (18). Node data (i.e. gene sets) are provided in Supplementary Material 1.

### Network reconstruction

Curated physical and genetic interactions documented for *Saccharomyces cerevisiae* were retrieved from BIOGRID (19) on 22/02/2017 (release 3.4.145). A dataset by Jansen *et al*. (20), where the authors used 10 permanent complexes as a basis for the classification of most *S. cerevisiae* complexes as permanent or transient, based on correlations with gene expression was used to identify the permanent and transient physical interactions. Dosage rescue, synthetic rescue, phenotypic suppression and positive genetic interactions were classified as “alleviating” responses, whereas synthetic growth defect, phenotypic enhancement, synthetic lethality, dosage growth defect, dosage lethality, synthetic haploinsufficiency, and negative genetic interactions were classified as “aggravating” phenotypic responses in the analysis of genetic interaction networks. Physical and genetic interactions were both regarded as bidirectional and the graph edges were of uniform weight. Multiple instances of experimental evidence for a single interacting pair from one or more independent resources were scaled down to a single documented instance. A genetically interacting pair was double-counted only when evidence for both aggravating and alleviating effects on the phenotype were documented. Self-loops (e.g. homodimers) were excluded from the analyses. Transcriptional regulatory network (TRN) interactions were accessed from YEASTRACT (21) (access date 22/02/2017). Transcription factor targets were used only when both DNA binding and expression evidence were documented. Transcription factors that do not have any curated targets were excluded from analysis. Regulatory evidence under all reported environmental conditions was included. Edge (interaction) data are available in Supplementary Material 1.

### Network and data analysis

Cytoscape v3.5.0 was employed for the removal of duplicate interaction pairs and self-loops, and for topological network analysis (22). Statistical analyses and network randomisation were conducted in MATLAB R2016b. Hubs were defined as nodes connected to at least 10% of the other nodes in the network and degree centrality was adopted as the hub-metric. Out-degree was employed as the characteristic parameter for the identification of principal transcription factors; i.e. the hub-like nodes in the bi-partite graph of the transcriptional regulatory network. Networks were normalised against the total number of nodes that they contained as reference. Dissimilarities in structural properties that could not be explained by the size effect were then determined by a “difference” metric, where a numerical network property was identified to be different from those for other networks only if the following relationship held true:

For any numerically representable network property *a_i_* for any network i = 1…n and where *j* ≠ *i*

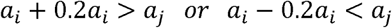

indicating that a property must differ by at least 20% from its peers in order to be identified as distinctly different.

A hypergeometric distribution was used to evaluate the statistical significance of functional enrichments (23). Princeton GO tools were employed for Gene Ontology Enrichment Analysis (http://go.princeton.edu/cgi-bin/GOTermFinder) accessed on (19/05/2017) (24). MeV: MultiExperiment Viewer was used for integrative data analysis (25). K-means clustering was employed for data classification (K = 2, distance metric = Pearson correlation, maximum number of instances to test all elements in the data set for cluster fit = 10000). Standard protocols were followed for multiple linear regression (MLR) and the determination of data normality (26). Symbolic regression (SR) was employed as described by (27,28). The gene expression data (Gene Expression Omnibus accession number GSE41094) was adopted from Rodríguez-Lombardero *et al.* (29). Rank correlations were employed in evaluating the emPAI values that correspond to protein abundance (30).

### Cell culture and pre-treatment for sub-cellular fractionation

The *S. cerevisiae* strain used in this study was BY4741 harbouring GFP-tagged Hsp82p (*MAT***a** *HSP82-GFP::HIS3 his3Δ1 leu2Δ0 met1SΔO ura3Δ0*) (31). Two separate cultures were grown to an OD_600_ of 0. 6 at 30°C in SD-His media with shaking. In total, 120 OD units were used for the nuclear preparation and 240 OD units were used for the chromatin preparation.

Pre-treatment of cells prior to lysis was based on (32). Cells were incubated at 5 OD units per mL of 25 mM Tris-HCl, pH 7.5, 10 mM TCEP for 10 minutes at room temperature. Cells were subsequently treated for 10 minutes at 30°C with 1 μg zymolyase 100-T (Nacalai Tesque) per OD unit of yeast, in spheroplasting medium (SD-His supplemented with 25 mM Tris-HCl, pH 7.5 and 1.2 M sorbitol), at 20 OD units per mL. Spheroplasts were washed once in SD-His, supplemented with 1.2 M sorbitol, at 5 OD units per mL.

### Sub-cellular fractionation

Nuclei were prepared according to (33), with the modification that lysis was carried out at 20 OD units/mL lysis buffer. This yielded a supernatant (S_NUC_) and a nuclear pellet (P_NUC_).

Chromatin was prepared according to (34), but in the absence of thiodiglycol. This yielded a supernatant (S_CHROMA_) and chromatin pellet (P_CHROMA_). The pellet was additionally treated with 500 U benzonase nuclease (Merck Millipore) for 30 minutes on ice and resolubilised in 30 mM Tris-HCl, pH 6.8, 2% (w/v) SDS.

### Sample preparation for mass spectrometry

Protein concentrations were estimated and aliquots representing 75 μg total protein per sample were resolved on 4-15% linear gradient polyacrylamide gels (Bio-Rad), which were then stained with Coomassie brilliant blue. Gel lanes were cut into equally sized bands (16 bands for S_50_ and P_50_; 8 bands for S_16_ and P_16_). Bands were destained, reduced (using dithiothreitol), alkylated (with iodoacetamide) and subjected to tryptic digestion at 37°C for 16 hours.

### Mass spectrometric analysis of samples

In all cases, approximately 1 μg of sample was injected on-column per mass spectrometry run. The tryptic digests from bands corresponding to S_50_ and P_50_ were analysed using a Dionex Ultimate 3000 RSLCnano UPLC (Thermo Fisher Scientific) system coupled in-line to a Q Exactive Hybrid Quadrupole-Orbitrap mass spectrometer (Thermo Fisher Scientific). The mass spectrometry method used was the same as described in (35), but with the modification that fragmentation was carried out on the twenty most intense ions per survey scan with charge state of 2+ and above.

The tryptic digests from bands corresponding to S_16_ and P_16_ were analysed using a nanoAcquity UPLC system (Waters) coupled in-line to an LTQ Orbitrap Velos Hybrid Ion Trap mass spectrometer (Thermo Fisher Scientific). The mass spectrometry method used was the same as described in (36), but the Orbitrap was operated at a resolution of 30,000 and fragmentation was carried out on the twenty most intense ions per survey scan with charge state of 2+ and above. The details of the proteomics analysis are provided in Supplementary Material 2.

### Mass spectrometry data processing

Files of raw mass spectrometry data were converted to Mascot Generic Format (MGF) using MSConvert (version 3.0.9283, Proteowizard). MGF files were searched using an in-house Mascot server (version 2.6.0, Matrix Science) against a canonical *S. cerevisiae* database, downloaded from UniProt (March 2017; 6,749 sequences). Samples corresponding to each gel lane were merged for searching, yielding a “master” results file of mass spectrometry runs for a given sample. Carbamidomethylation of cysteine was specified as a fixed modification and oxidation of methionine as a variable modification. A 1% FDR threshold for protein identifications was imposed, based on the search of a decoy database of reversed protein entries. Relative protein quantification was inferred using the built-in emPAI score (30) feature of Mascot (Supplementary Material 3).

## Results and Discussion

We investigated the network organization and transcriptional regulation of the genes and their products involved in “Protein Transactions” (PT). We define PT as comprising the biosynthesis, modification, localization, and degradation of proteins. This set of processes is orchestrated by 2533 gene products in the yeast *Saccharomyces cerevisiae*, approximately 38% of the genes encoding protein, rRNA, snRNA, snoRNA or tRNA molecules. The physical, genetic and transcriptional regulatory interaction networks of this set of gene and gene products were constructed and these networks were topologically investigated to identify whether network topology could provide biological insight, which we will refer to as ‘functional signatures’ from this point forward. We investigated the effects of both the size and functional homogeneity of the network on the detection of these functional signatures. The ubiquity of the identified signatures was then assessed by analysing other large networks of gene sets dedicated to other specific functions in yeast; gene expression (GE, 2128 gene products), cell cycle (CC, 786 gene products), and small molecule metabolism (SMM, 835 gene products). A summary of the biological networks constructed for all sub-systems and for the genome-scale system of 6447 genes and their products are provided in Supplementary Material 4 and in Figure 1a.

**Figure 1.**
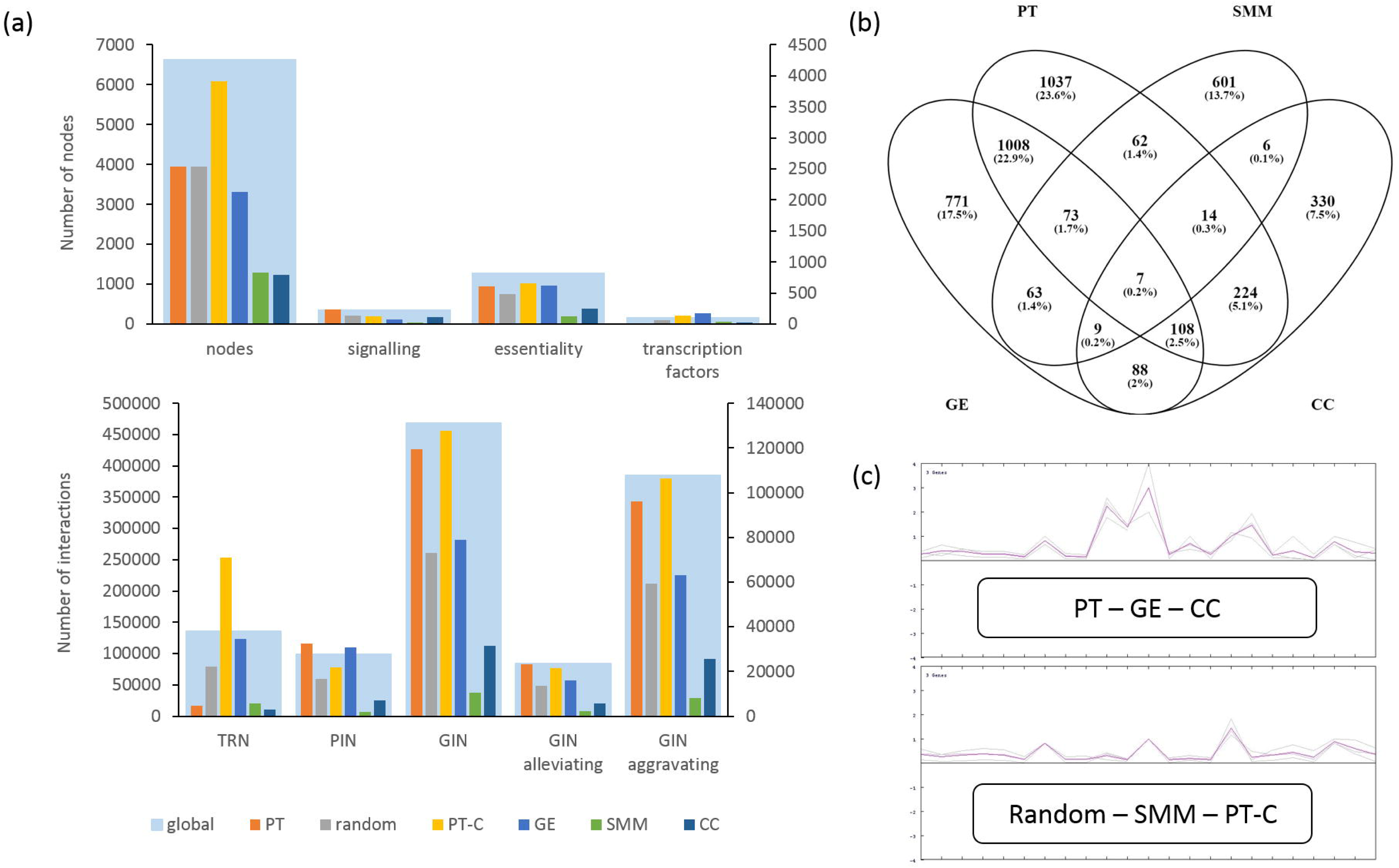
Summary of network properties of the systems under investigation, (a) The number of nodes and interactions for each system under investigation; the global network, PT, the random network, PT-C, GE, CC, and SMM are provided in two plots. In both plots, the sub-systems are compared against the global network denoted by the light blue bar in the background. The primary vertical axis on the left hand side represents the values for the global network; whereas the secondary vertical axis on the right is used for the sub-systems, (b) The overlap of nodes amongst different sub-systems with defined biological functionality. The percentage values given in parentheses denote the relative proportion of the numbers among the total number of unique nodes from all 4 sub-systems, (c) Classification of the sub-networks by K-means clustering based on the topological properties summarised in Table SI. The two main classes of sub-networks are highlighted in the rounded boxes on their respective similarity profiles. Note that the abbreviations adopted in the Figure are consistent throughout the article.

We discuss here our findings on the necessary and sufficient criteria for gaining biological insight by topological analysis of the protein transactions in yeast. This analysis revealed unusual properties regarding transcriptional regulation.

### Transcriptional regulation of protein transactions in yeast

The topological analysis (Supplementary Material 4) of the transcriptional regulatory network (TRN) of PT indicated that only 17 of the 2533 gene products in the yeast protein transactions network encoded TFs. Transcription factors in the protein transaction network indicated either a direct role in the regulation of the protein transaction by transcription, or regulation of a transcription factor by a protein transaction.

The number of genetic and physical interactions reported between these TFs and the protein transactions domain were significantly lower than what would be observed for all transcription factors in the global network (under-enrichment by 2.98-fold with *p* — *value* < 10^−189^, and by 2.78-fold with *p* — *value* < 10^−101^, for PIN and GIN, respectively). However, an investigation of these 17 TFs in the global network revealed that they had a significantly higher number of interactions than those for all TFs (over-enrichment by 1.89-fold with *p* — *value* < 0.002, and by 1. 77-fold with *p* — *value* < 0.004, for PIN and GIN, respectively). This indicated that although these TFs were highly interactive nodes, their specific role in protein transactions was tightly defined and allowed less interaction (i.e. communication) with other nodes in the network. Despite the fact that none of these TFs was identified as a hub in the PIN and GIN of protein transactions, they were all linker proteins between hubs in both networks. In other words, despite their relatively low degree, their “strength” (37), indicating how important or central they are for the network, was very high.

The non-TF nodes of the TRN of PT were observed to be significantly enriched for genes that can complement human gene function (11% vs 8.5% of that of the global TRN, 1.31-fold over-enrichment with *p* — *value* < 10^−09^), reflecting the conservation of genes involved in protein transactions across the eukarya. Hsf1p, a heat shock protein regulating gene transcription in response to stress, was the only transcription factor that is included in the TRNs of both PT and the global set, and which has been experimentally shown to complement the function of its cognate human gene. In contrast, 42 transcription factors were present in SMM TRN. Therefore a much smaller repertoire of transcription factors were associated with the large PT network (2533 nodes) compared with the SMM (835 nodes) or the global network (6447 nodes).

### Hubs in the PT network

Seven genes (*VAM6, RPN11, HSP82, CDC28, UBP3, RPN10* and *BRE5*) were identified as hubs in both the physical and the genetic interaction networks of PT, whereas no common hubs were identified for the PIN and GIN of the global set. These genes were tightly connected (through physical or genetic interactions) with other members of PT. Since these seven genes could only be defined as hubs for the PIN and GIN of PT, they are thought to assume a pivotal role in protein transactions, whereas their impact on the global network was less dominant. The genetic interactions involving these common PI/GI hubs was found to be exclusively aggravating. Indeed, synthetic-lethal genetic interactions were previously reported to significantly overlap with protein–protein interactions, and were reported to be largely confined to genes within pathways that contain at least one essential gene (38). The hubs of the PT GIN (and consequently the aggravating GIN) were significantly enriched for the GO Process Terms ‘protein localisation’ (p — *value* < 10^−32^) and ‘transport’ (*p* — *value* < 10^−30^). A similar analysis on the hubs of the PT PIN revealed no enrichment other than the uninformative common parents of the PT network terms. In contrast, the hubs in the physical or genetic interaction networks of the global set were not significantly enriched for any biological process.

The identification of hubs found in both the physical and genetic interaction networks for the PT gene/gene product set led us to search for common genes that act as principal TFs in the TRN and as hubs in either the PIN or GIN of PT. However, the nodes of the TRN were very significantly underrepresented in either the GIN or PIN of PT in comparison to what was observed for the global set (under-enrichment by <2.5-fold; *p* — *value* < 10^−121^, and by 3.5-fold; *p* — *value* < 10^−551^ compared to expectation, respectively). This could potentially be the result of the downsizing of the PT TRN, indicated by the very low number of TFs identified within the PT subset.

### Small network size is neither necessary nor sufficient

Identification of the features of the PT interaction networks that were different from those of the global interaction networks of yeast raised two related questions regarding the topological analysis of biological systems: (i) Did these system-specific characteristics appear due to the fact that the global network was reduced down to 38% of its original size, in line with earlier arguments on the necessity to work with smaller networks in order to extract biologically relevant information (39), or (ii) did functional coherence among the gene members contribute to the understanding of the system under investigation? We addressed these questions by investigating 1000 randomly selected sub-sets of the global yeast gene set that comprised 2533 genes (the size of the PT set). On average, 39.30 ± 0.79% of these random sets were associated with genes from the PT set. Since, the PT set itself constituted 38% of the global set in yeast, we concluded that this randomisation was both a robust and realistic downsizing of the global set. One of these 1000 random sub-sets was then randomly selected to construct genetic interaction, protein interaction, and transcription regulatory networks. The characteristics of these networks were compared to those of both global gene/gene product set and the *bona fide* PT gene/gene product set (Table 1). Node-wise comparisons revealed that the random network was a realistic scale-down of the global network whereas PT was enriched with signalling-associated or essential genes. Although both the random and the PT subset had a higher number of nodes identified as hubs, this was considerably more prominent in the physical and genetic interaction networks of PT, and was only a slight increase for the random subset.

**Table 1.** A network-based comparison of the protein transactions *gene set* against a random *set of the same size*

|  | *PT*<br>*global* | *random*<br>*global* |
| --- | --- | --- |
| *fraction of genes contributing to network analysis* | 0.38 | 0.38 |
| *fraction of signalling-associated genes* | 0.65 | 0.36 |
| *fraction of essential genes represented* | 0.47 | 0.37 |
| *fraction of hubs in PIN* | 1.57 | 0.50 |
| *fraction of hubs in GIN* | 2.39 | 0.42 |
| *fraction of TFs* | 0.09 | 0.37 |
| *fraction of principal TFs by outdegree* | 0.11 | 0.46 |
| *fraction of interactions in TRN* | 0.03 | 0.16 |
| *fraction of interactions in PIN* | 0.33 | 0.17 |
| *fraction of interactions in permanent protein complexes* | 0.64 | 0.16 |
| *fraction of interactions in transient protein complexes* | 0.33 | 0.17 |
| *fraction of interactions in GIN* | 0.25 | 0.16 |
| *fraction of alleviating genetic interactions* | 0.28 | 0.16 |
| *fraction of aggravating genetic interactions* | 0.25 | 0.15 |

Scaling down reduced the available information at the interaction level for both random and PT networks since only the interactions between the genes of a given set were considered in the analysis. The random GIN and PIN were observed to be more severely affected, having fewer interactions, than PT GIN and PIN. The functional homogeneity in PT resulted in a higher within-graph connectivity characterised by higher network density by 45% and 40%, and shortened the characteristic path lengths by 10% and 8% for PIN and GIN, respectively. However, the transcriptional regulatory events, at both the node and interaction levels, were severely underrepresented in PT – indicating that there were (i) fewer nodes identified as TFs, (ii) fewer nodes identified as target genes of these TFs, and (iii) fewer TF-gene binding interactions between (i) and (ii). This feature could not be explained by differences in any topological measures of the TRN, and therefore was regarded as novel biological insight, rather than an inherent artefact of topological network analysis.

This analysis of random networks demonstrated that reducing the size of the set of nodes from which the biological networks are constructed did not markedly affect the analysis of the network; rather, it was found that functional integrity within the set allowed novel biological insights to be garnered from the available information.

### Functional specificity is necessary but not sufficient

The next question to be explored was whether topological features identified for PT were peculiar to the protein transactions gene/gene product set or whether they could be found for other well-defined gene/gene product sets. For this purpose, four functionally distinct (but non-exclusive) subsystems: PT, GE, CC, and SMM were analysed. Although each subsystem constituted all the genes/products attributed to a broad but single cellular process, many genes had multiple functionality, and were thus shared across the different subsystems. The overlap between PT and GE was especially striking, with nearly 47% of all PT shared with GE (Figure 1b), largely due to the presence of the biological process “translation” in both protein transactions and gene expression. Although each subsystem had unique members, it was not possible to identify a biologically relevant functional subgroup that would exclude overlapping genes from the analysis, and still provide meaningful large datasets.

Since the number of genes in each functional subset was different, the topological features for the PINS, GINS and TRNS of PT, GE, CC, and SMM were normalised for their number of constituent nodes relative to that for the global network (Table 2, Supplementary Materials 4, 5). A broad classification of these topological features by k-means clustering revealed that the topological network properties of PT, GE and CC were similar, and clustered separately from those of the random subsystem, SMM and PT-C, which were also similar(Figure 1c). Interestingly, the network topology of SMM was different to that of the other functional networks. The gene products of the members of SMM are most frequently involved in metabolic routes in the cell or in the regulatory events controlling these metabolic pathways. The flow and exchange of information often takes place through the small molecules, themselves, rather than the genes or gene products in the metabolic network. In contrast, the expression of genes or proteins, or the cell cycle, where the genes or gene products themselves undergo changes, relay or exchange information, and this could have resulted in the topological properties of the SMM being different to those of PT, GE and CC. Parallel linear cascades of information flow with fewer connections are more common in metabolic networks. This functional difference was observable at the level of network connectivity. The PIN diameter of SMM was identified to be the largest (d=11) among all PINs, and 37.5% higher than the second largest identified in this study. In line with these results, Szappanos *et al.* reported a unique property of metabolic networks, highlighting the underrepresentation of genetic interactions, specifically of those that take place between pathways, in yeast (40).

**Table 2.**
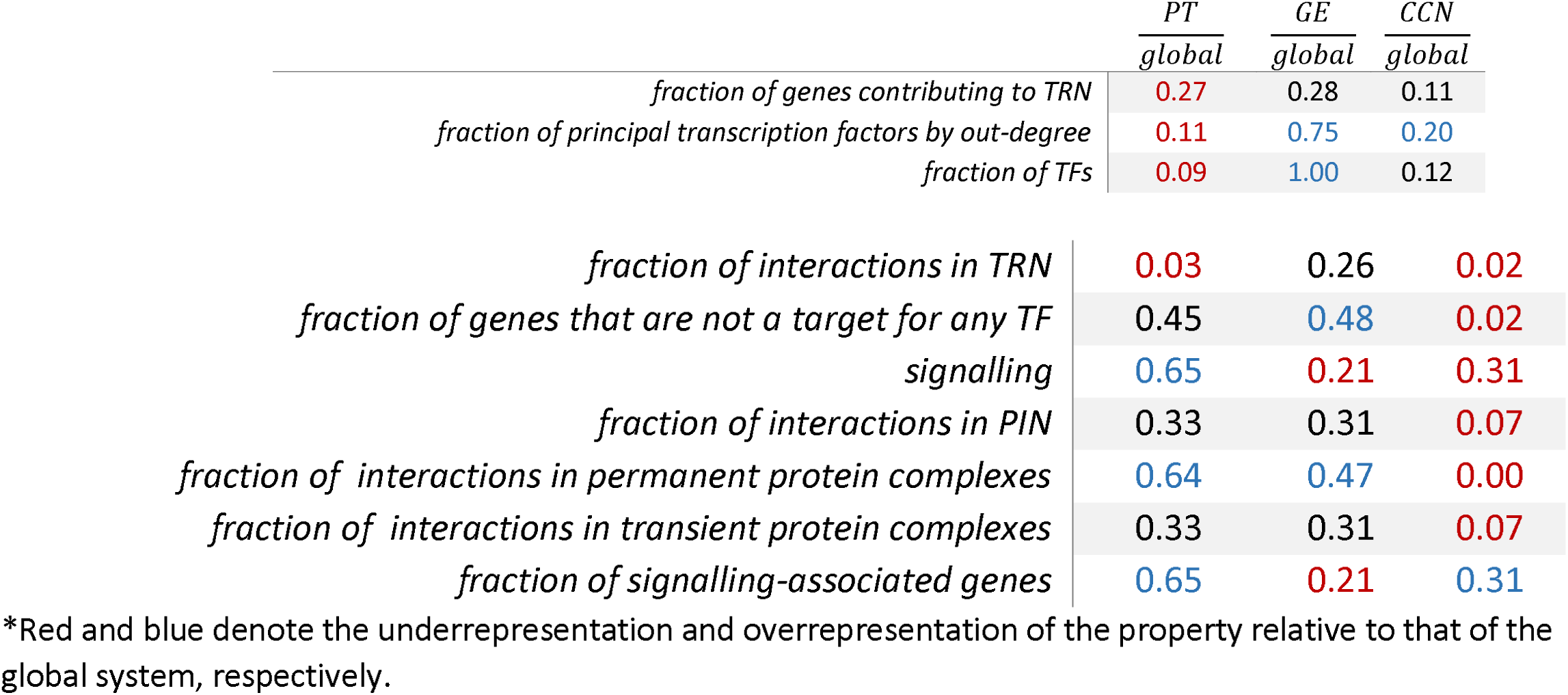
Distinct network properties of PT, GE, and CC that cannot be explained by a size effect.*

|  | *PT*<br>*global* | *GE*<br>*global* | *CCN*<br>*global* |
| --- | --- | --- | --- |
| *fraction of genes contributing to TRN* | 0.27 | 0.28 | 0.11 |
| *fraction of principal transcription factors by out-degree* | 0.11 | 0.75 | 0.20 |
| *fraction of TFs* | 0.09 | 1.00 | 0.12 |
| *fraction of interactions in TRN* | 0.03 | 0.26 | 0.02 |
| *fraction of genes that are not a target for any TF* | 0.45 | 0.48 | 0.02 |
| *signalling* | 0.65 | 0.21 | 0.31 |
| *fraction of interactions in PIN* | 0.33 | 0.31 | 0.07 |
| *fraction of interactions in permanent protein complexes* | 0.64 | 0.47 | 0.00 |
| *fraction of interactions in transient protein complexes* | 0.33 | 0.31 | 0.07 |
| *fraction of signalling-associated genes* | 0.65 | 0.21 | 0.31 |

\*Red and blue denote the underrepresentation and overrepresentation of the property relative to that of the global system, respectively.

Networks of all functionally coherent sets were comprised of fewer interactions, which could not be restored by normalising the networks with respect to the number of nodes that possess. This downsizing effect was considered as an artefact of the network reconstruction process, due to only interactions within the members of each set being considered, and is not discussed further. On the other hand, the GE and SMM networks were enriched for the number of TFs that are present among the participating genes/products; this contrasts with PT, in which TFs were strikingly underrepresented. This emphasises the significance of the identification of substantially fewer transcriptional regulatory interactions among the members of PT. Consequently, fewer nodes were identified as principal TFs (by out degree) in the TRN of PT than of GE, SMM, or CC. On the other hand, more hubs were identified in the GINs and PINs of all functionally specific sets, with CC networks possessing more hubs than the others (Supplementary Materials 4, Figure 2).

**Figure 2.**
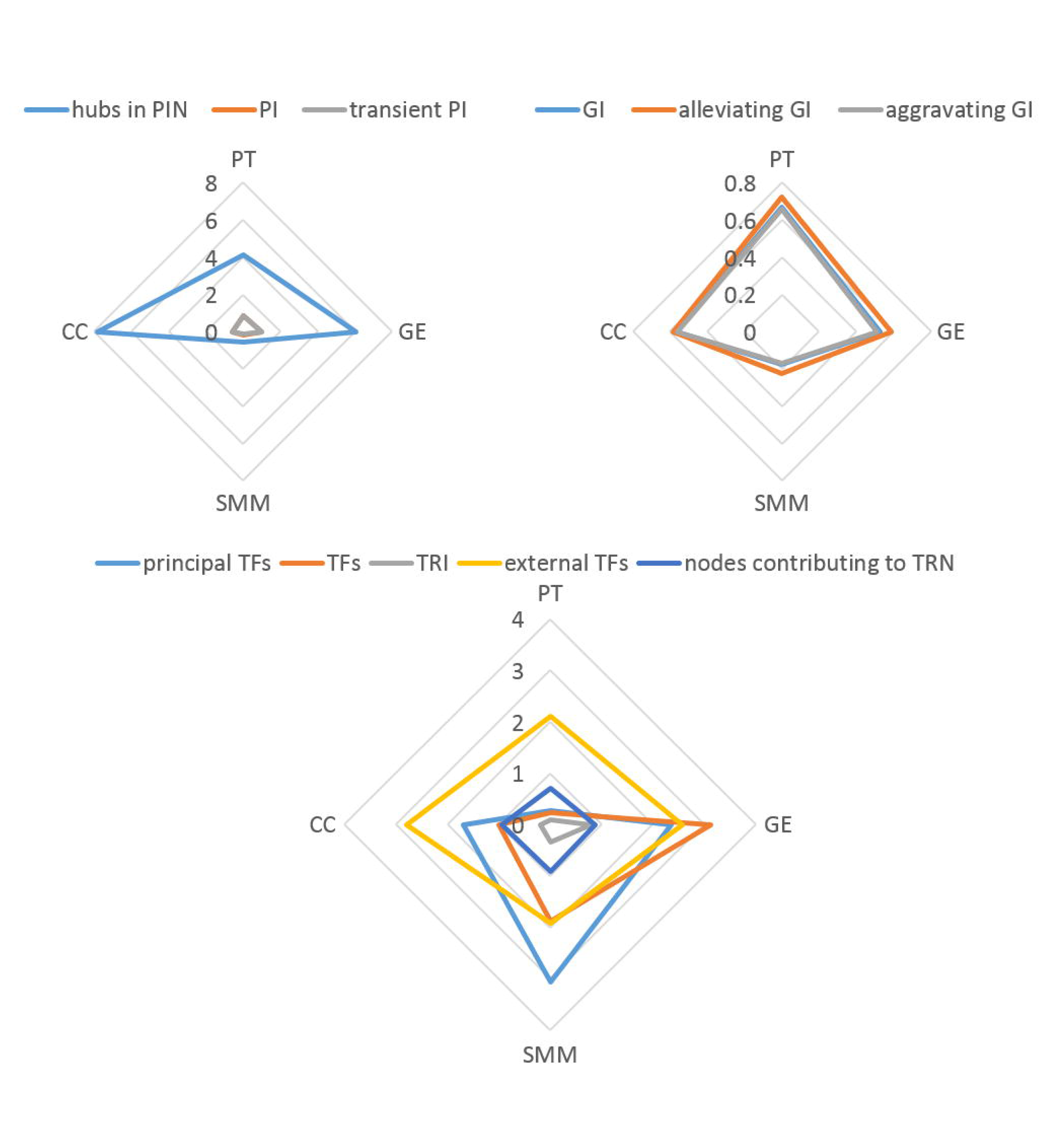
Relative differences in topological parameters about functionally defined subsystems. Radar plots were employed to highlight the relative differences between PT, GE, SMM, and CC. The size of each subnetwork is normalised against that of the global network and the topological properties displayed in the Figure represent the ratio of the numerical value for that property to that of the global network.

The underrepresentation of TFs within the PT set was striking, but excluded the possibility of any non-PT TF regulating the transcription of PT genes. This was addressed by identifying all TFs, which had documented binding and expression evidence for the members of PT, GE, SMM, or CC. A size-normalised analysis indicated 1-2 fold additional TFs regulating the transcription of the genes of these functional sets but which were not themselves members of these sets, but no significant differences were observed between the sets (p-value>0.1) indicating that the transcriptional regulatory signature of PT remained unique among the analysed sets (Figure 2).

We investigated the underrepresentation of TFs in the PT TRN further by artificially replacing a fraction of the PT nodes by those that were known not to have any role in protein transactions (i.e. introducing false positives). Even upon the random replacement of only 10% of the nodes, the reconstructed TRN had up to 75% more transcription factors and 63% more interactions. If 40% of the nodes were randomly replaced by those that are known not to be involved with protein transactions, the reconstructed TRN had 237% more TFs and 257% more interactions. Despite extensive studies, a considerable fraction (nearly 17%) of the protein-coding genes in *S. cerevisiae* are still not assigned to known biological processes (18). Therefore, such nodes would highly likely be encountered, or even be included unintentionally, in network analysis. As an additional exercise, we tested if the inclusion of such nodes would interfere with the topological structure of the TRN. We introduced nodes with unknown functionality so as to increase the size of PT by 10%. This expanded the PT-TRN by increasing the number of TFs and transcriptional regulatory interactions by only 13% and 11%, respectively. Similarly, a 40% increase in node size was accompanied by a 39% increase in the number of TFs in the TRN and 49% more transcription factor-gene interactions to be introduced into the PT-TRN. Therefore, the expansion of the PT network by unknown nodes was observed to be correlated with the fraction of nodes introduced and did not create any structural changes.

The reorganisation of the hub and principal node structures in these genetic, physical, and transcriptional regulatory interaction networks was examined for any unique signatures that might provide further biological insight. All hubs in the PIN of SMM (vs 14% of the hubs in the global network) and 50% of all hubs in alleviating GIN of CC (vs none in the global network) were observed to be yeast genes that are able to complement the cognate human gene function. More than twice as many PT, GE, and CC PIN hubs were essential genes or genes whose absence impairs growth significantly, in comparison to the global PIN hubs (45%, 48%, and 52%, respectively, vs 21%). In contrast SMM GIN hubs comprised markedly fewer such genes (8% vs 52% of that of global GIN). The principal TFs identified for PT were strikingly different, being both enriched for growth and viability genes (43% vs 15% for the global TRN) and for functional orthologs of human genes (14% vs 1% for the global TRN) (Supplementary Material 5).

Characteristic signatures were determined in the physical and genetic interaction networks of CC and SMM, and in the transcriptional regulatory network of PT. A striking result was that a unique topological signature could not be attributed to the GE for any of the network parameters studied despite the set’s substantial overlap with PT. This indicated that topological signatures were indeed function-specific rather than ubiquitous in the present analysis.

### Functional homogeneity is necessary

The final question we addressed was whether the biological relevance of our findings could be attributed to the fact that the network analysis was carried out on a functionally coherent set, or whether the topological network analysis of a set of gene products that can be attributed to different and unrelated biological processes could also be as informative. For this purpose, a similar analysis to that conducted for PT was conducted for PT-C. The PT-C set was functionally heterogeneous; it comprised gene products from various specific functional subnetworks, some of which were investigated here; a major fraction of this subset belonged to the small molecule metabolism. The topological properties of the interaction networks for the PT-C gene/gene product set closely resembled that of the random networks. Transient protein-protein interactions and aggravating genetic interactions were observed slightly more in the PT-C subset than among the members of the random network, but this difference was not statistically significant. Both the PIN and GIN were sparser than randomly constructed networks of the same number of nodes, by 50% and 28%, and had longer characteristic path lengths by 17% and 8%, respectively. Only 2 hubs were identified in the PIN and 11 in the GIN of PT-C. These findings indicated that PT-C did not reveal any further biological insight into the system, possibly due to this weak connectivity between network components (nodes) since these nodes represented a collection of genes/products with diverse functions in the cell, and did not constitute a cohesive subset (Supplementary Material 4).

### Further insight into transcriptional regulation of protein transactions

PT, CC and GE, which were clustered together with respect to the topological properties of their cellular networks (Figure 1c), were investigated for the subcellular localisation of their proteins, using existing GO annotation. The gene products of CC were significantly localised only in the nucleus (*p* — *value* < 10^−90^) whereas those of GE significantly localise in the nucleus, or in the cytosol (*p* — *value* < 10^−80^). On the other hand, the nucleus (or any of its sub-compartments) were not among the enriched cellular compartments for the gene products of PT (Figure 3). The over-representation of the PT gene products in the nuclear compartment related only to the nuclear outer membrane-endoplasmic reticulum membrane network term, only. This difference in localisation pattern, along with the underrepresentation of the transcriptional regulatory elements, led to the investigation of the proteomic landscape of yeast focusing on the nuclear-enriched and, more specifically, the chromatin-enriched fractions of the yeast cell, as well its remaining subcellular compartments (Supplementary Material 3).

**Figure 3.**
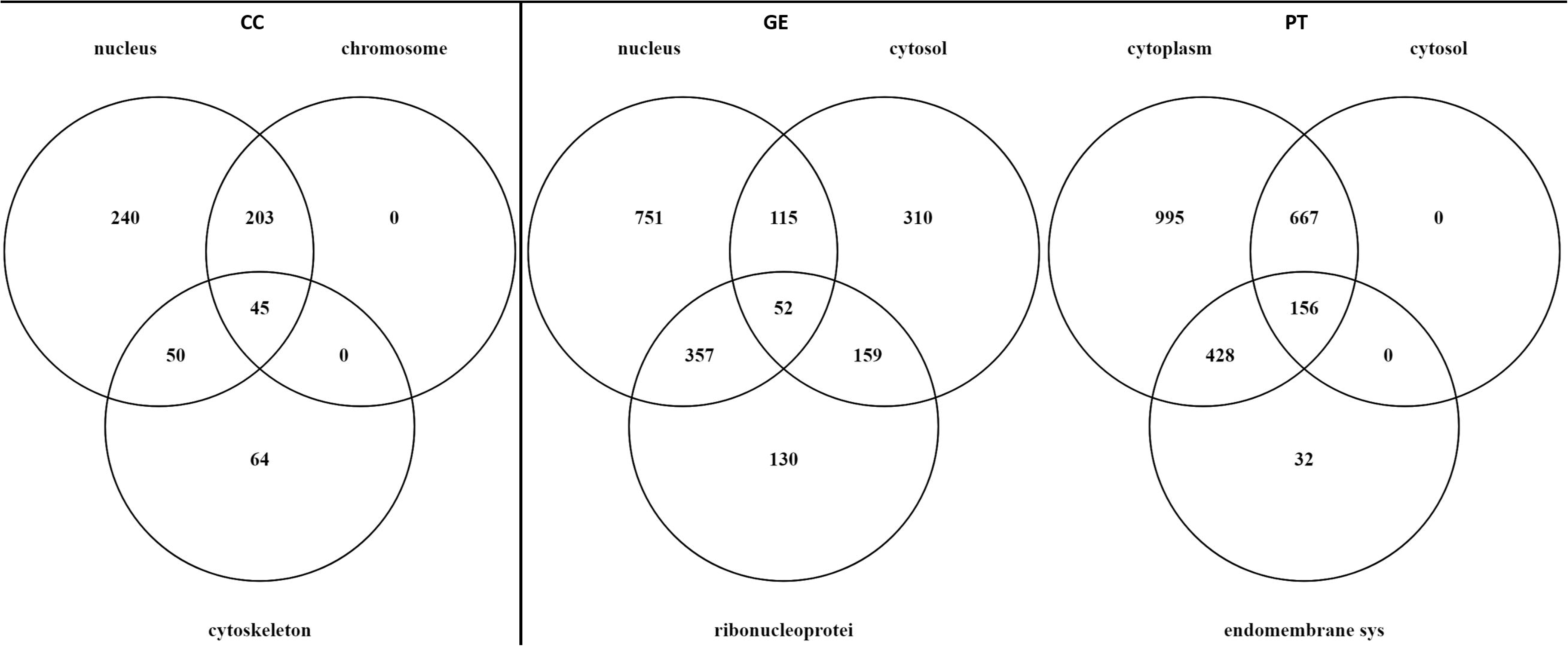
Principal subcellular locations for the genes in the subnetworks of the PT-GE-CC class. The main subcellular components were identified by the Gene Ontology Component and only components populated by at least 30% of the nodes constituting a subsystem are displayed. NB: Cytoplasm, as defined by the Gene Ontology, is comprised of all of the contents of a cell excluding the plasma membrane and nucleus, but including other subcellular structures. Cytosol, on the other hand, defines the part of the cytoplasm that does not contain organelles, but which does contain other particulate matter, such as protein complexes.

A total of 1572 and 3272 proteins were identified in the analysis of the chromatin-enriched and nuclear-enriched subcellular fractions, respectively. In order to simplify the description, we will refer to these two independent experiments as CHROMA and NUCLEA, respectively. 1541 of the 1572 proteins identified in CHROMA were also identified in NUCLEA. Proteins, which were exclusively detected in the chromatin-enriched fraction, were significantly enriched for involvement in chromatin-remodelling processes, as would be expected (*p* — *value* < 10^−9^), and those exclusively identified in the non-chromatin fraction were significantly associated with small molecule, oxidation-reduction, and nitrogen compound metabolic processes (*p* — *value* < 10^−7^). Proteins specific to the nuclear-enriched fraction were associated with the cellular component organisation or biogenesis process term (*p* — *value* < 10^−31^). In contrast, proteins detected exclusively in the supernatant of NUCLEA were enriched for small molecule metabolic process (*p* — *value* < 10^−18^). Approximately 80% of the proteins identified in CHROMA had a lower abundance in the chromatin-enriched subcellular fraction than in the supernatant, and they were identified to be functionally associated with small molecule metabolism (*p* — *value* < 10^−70^). In contrast, 76% of the proteome identified in NUCLEA were present at higher levels in the nuclear-enriched subcellular fraction than in the supernatant (cellular component organization or biogenesis (*p* — *value* < 10^−112^)), possibly indicating active gene expression process ongoing in the non-chromatin nuclear-enriched subcellular compartment. Our analysis of the shotgun proteome of the nucleus (specifically, that of the non-chromatin fraction) indicated its high specificity, with only a few proteins ubiquitously being detected in more than one enriched fraction (Supplementary Material 3).

The proteomes identified in CHROMA and NUCLEA were both significantly enriched for the components of the PT TRN (*p* — *value* < 10^−13^) indicating satisfactory coverage of the investigation space by the experimental datasets. Only 7 of the 17 TFs in the TRN of PT were detected in the proteome analysis; Gzf3p, Hsflp, Raplp, Rlmlp, Rtg3p, Skn7p, and Mot2p. However, neither the chromatin-enriched nor the nuclear-enriched proteome was significantly overrepresented for the 17 TFs of the PT set (*p* — *value* > 0.1). Furthermore, the TRN components of PT were underrepresented among those proteins identified in the chromatin-enriched subcellular fraction, as well as in both fractions in NUCLEA data, with a low level of significance associated with the findings (*p* — *value* < 0.3). On the other hand, the TRN components were significantly overrepresented in the supernatant of the CHROMA data, which is enriched with the non-chromatin fraction of the nuclear components (*p* — *value* < 10^−4^), possibly indicating a role in the regulation of nuclear processes by protein transactions (Supplementary Material 3).

Sixteen genes in PT were identified as targets for all of the 7 TFs detected in the proteome analysis of the nucleus. These TFs and the genes that are regulated by all 7 TFs, retrieved from YEASTRACT based on experimental and expression evidence, were used to construct a matrix of TF protein abundance (obtained from the nuclear-enriched fraction of the NUCLEA dataset) against the gene transcript abundance (obtained from (41) under similar experimental conditions to those for the proteomics analysis carried out here). This information matrix was then employed to construct a mathematical abstraction of the functional fingerprints of these transcriptional regulatory proteins of the yeast protein transactions (Figure 4a). The non-normality of the population from which the data matrix was constructed (fat-tailed, with excess kurtosis = 2.4, positively skewed = 1.5) was taken into consideration in constructing a non-linear regression model employing symbolic regression, and this method was observed to model the data with a satisfactory goodness-of-fit (*R*^2^ = 0.82) (vs. MLR *R*^2^ = 0.66) (Figure 4b, Supplementary Material 3). This empirical mathematical model allowed us to predict the transcription of 16 common targets of these 7 TFs with reasonable accuracy. Protein transactions are closely related with the growth and proliferation of the cell. Growth-associated processes were previously reported to depend on precise transcriptional control (42), and this reasonably high level of success in the prediction of the expression levels of these 16 transcripts could be attributed to the functional specificity of the PT network. We note that this analysis rests on the principles for constructing non-linear predictive models of gene expression and the performance of these models is limited by the size of the dataset under investigation. Therefore, the small size of the dataset investigated here should be taken into consideration in evaluation of the results.

**Figure 4.**
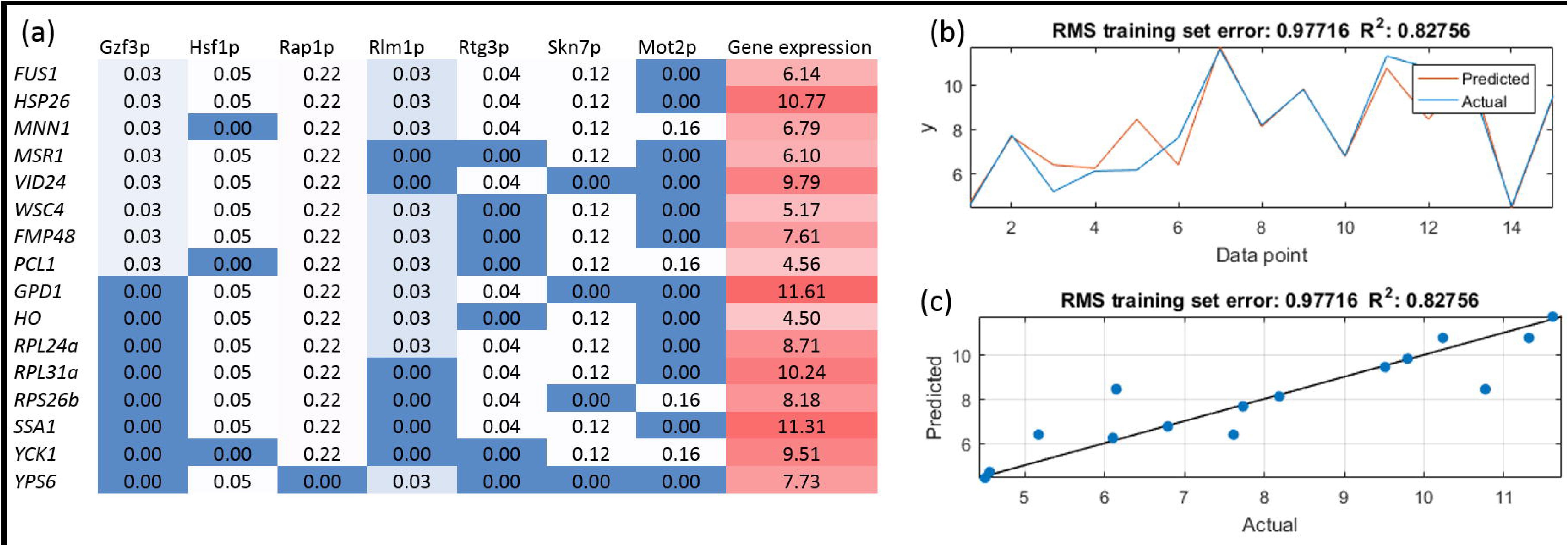
Transcription factor abundance as a proxy for target gene mRNA expression, (a) Nuclear abundance patterns of the transcription factors from PT quantified in the proteomics study. The mRNA levels for their target genes were adopted from a study conducted under similar conditions (41). (b) and (c) The goodness of fit measures for the non-linear model predicting target gene mRNA expression levels utilising transcription factor protein abundance.

## Conclusions

We employed simple network analysis to gain novel biological insights about the extensively studied protein transactions in yeast. Our analysis showed that despite the high number of genes involved in these protein transactions, which comprised of nearly 38% of the gene products of *Saccharomyces cerevisiae*, only a limited number of transcription factors were associated with this dataset, and through a small network of interactions. Transcription factors are only annotated to the regulation of a process, if that process is directly initiated/activated by that target gene’s transcription, and not if the process is activated by an independent mechanism following the biosynthesis of the participating gene products. Transcription factors annotated in the PT network were therefore i) directly regulating the process performed by the PT gene or ii) represent an instance where a PT regulates the transcription factor.

Although the TFs identified were highly interactive nodes, their specific role in protein transactions was tightly defined, and allowed less interaction (i.e. communication) with other nodes in the network. This reduced interactivity of the identified TFs makes it more likely that they represent instances where the protein transaction regulates a TF (see ii), above). Inspection of the TFs in the PT network showed that 9 identified protein transactions regulating transcription (usually *via* protein recruitment to chromatin; Gis1p, Gzf3p, Haplp, Hsflp, Maclp, Mot2p, Raplp, Rphlp, and Xbplp), and 3 represented TF’s directly regulating processes performed by the PT (Hac1p, Skn7p, and Yap1p; all related to the Unfolded Protein Response). The remaining 4; Rlm1p, Rpn4p, Rtg3p, and Smp1p were annotated in error. They were either annotated by computational inference (IMP-type annotation), or their function was not correctly described. For example, Rpn4p has a role in regulating the transcription of the proteasome, but is not involved in the positive regulation of proteasomal ubiquitin-dependent protein catabolic process, as previously annotated. These annotation errors were corrected in Saccharomyces Genome Database upon the authors’ request.

In contrast, 42 transcription factors were present in SMM TRN, all representing direct regulation of SMM processes by transcription. Therefore a much smaller repertoire of transcription factors were responsible for the direct regulation of processes associated with the PT network compared with the SMM or the global network. Even when considering TFs regulating PT genes, which were not themselves in the PT set, the network still had a reduced number of TFs indicating that the transcription of these genes is also orchestrated by a smaller set of TFs.

Our analysis was shown to be independent of the size of the networks that were constructed; large networks were shown to be as informative as smaller networks. On the other hand, the ability to capture biological information from network topology was shown to depend predominately on the functional homogeneity of the genes; networks that were constructed from gene components that play roles in a specific biological function were observed to be informative, whereas multiple functions in the network caused interruptions in connectivity. Moreover, different topological properties were identified to be specific to different biological processes.

This analysis has demonstrated that the “execution” of protein transactions is rarely directly transcriptionally regulated but the execution of metabolic processes often is. The SMM network provides the elemental building blocks required for energy generation and growth (nucleotides, amino acids, co-factors). Pathways in this network are often activated by direct and specific transcriptional activation in response to nutrient and environmental conditions. Gene products are often functional as expressed, and the metabolic pathway is fine-tuned by “within pathway” rate-limiting steps executed by substrate activation and inhibition when nutrients are replete or limited. The large number of transcription factors controlling the processes encoded by this network is a consequence of the need to respond rapidly and specifically to the external environment and express pathway components to utilise available nutrients.

In contrast the processes orchestrated by the PT network are frequently related to managing the temporal events of cell division once per cell cycle. Here, the effectors are provided transcriptionally, by a smaller number of transcription factors, constitutively or in advance of cell cycle’s phase-specific needs. Expression of a gene product does not appear to directly regulate these processes except in response to stresses; rather, they are largely activated post-translationally in response to signalling pathways. The observations we make here are consistent with the direct regulation of metabolic processes by transcription, and the regulation of cell division processes by post-translational modification.

Our observations were also supported by the identification of a relatively low number of transcription factors (7) in the nuclear-enriched fraction of the proteome, which were universally responsible for the transcriptional regulation of protein transactions in yeast under normal growth conditions. A model-based analysis on the TFs and their common target genes involved in protein transactions indicated that interestingly, the TFs that regulate the transcription of some candidate genes could be used to construct a model to describe the expression of their common target genes. Although it would not have been possible to predict the transcription levels of all target genes of a single TF, the transcription of a group of genes could be mathematically described by the protein expression levels of functionally linked TFs that target all these genes.

We feel that this notion of regulation should be approached carefully, as metabolic regulation information at the global scale is rarely available. So the “regulation” discussed in this context remains strictly “hierarchical” (43) restricted to epigenetic, transcriptional, translational or post-translational control. Protein transactions, because they are mainly relevant to large numbers of proteins (e.g. translation relating to all proteins), or proteins always required at least once every cell cycle, are not transcriptionally regulated; they are post-transcriptionally regulated and so they are controlled by a small number of transcription factors. Although this hypothesis will have its exceptions, what we observe is significant. The results imply a very strong functional difference, especially considering that all of “translation” is part of GE as well as PT.

## Data, code and materials

The datasets supporting this article have been uploaded as part of the supplementary material. The proteome data are available via ProteomeXchange with identifier PXD007525.

## Authors’ contributions

DD carried out the network and data analysis, designed the study and drafted the manuscript; DJHN carried out the proteomics analyses and drafted the manuscript; VW participated in the design of the study, supervised ontology based analysis, and drafted the manuscript; VW and SGO helped interpret the data, KSL and SGO conceived of the study, coordinated the study and helped draft the manuscript. All authors gave final approval for publication.

## Acknowledgements

The authors thank Professor Maya Schuldiner for the strain used in the study and Dr Mike Deery for mass spectrometric analysis of the proteomics samples.

## Supplementary material

### Supplementary Material 1

This document includes the node information, the genetic interaction network (GIN), physical interaction network (PIN), transcriptional regulatory network (TRN), and the permanent protein complexes for the global network, PT, the random network, the PT-C, GE, CCN, and SMM. The file also includes miscellaneous data on signalling-associated proteins, gene essentiality, human functional homology, haplosufficiency, and permanent protein complex information (format: .xlsx).

### Supplementary Material 2

This document details out the experimental protocols employed in the proteomics analysis. (format: .docx).

### Supplementary Material 3

This document includes the relative abundance (provided as emPAI scores) of the proteins identified in the mass spectrometric analysis of the chromatin enriched and nuclear enriched subcellular fractions along with the details on the statistical and model-based analyses of the datasets in the subsequent worksheets (format: .xlsx).

### Supplementary Material 4

Table SI summarises the topological analysis of the networks reconstructed in this study for of the systems under investigation; the global network, PT, the random network, PT-C, GE, CCN, and SMM (format: .docx).

### Supplementary Material 5

Table S2 summarises the analysis of the hubs in the genetic (alleviating and aggravating genetic interaction networks also investigated separately) and physical interaction networks as well as the principal TFs in the transcriptional regulatory network of the systems under investigation; the global network, PT, the random network, PT-C, GE, CCN, and SMM. The fraction of the hubs whose null mutants are reported to have impaired viability, and those that can complement human gene functionality are provided in the Table.

